## Supplementary Text for "Serostatus Testing & Dengue Vaccine Cost-Benefit Thresholds"

### CONTENTS


List of Tables 1

List of Figures 1

I. Assumptions 3

II. Cost-Benefit Equations 3
- A. Definitions 3
- B. Status Quo and Vaccination Only Costs ($C_0$, $C_V$) 4
- C. Model with Free Testing 5
- D. Cost-Benefit Constraint Equations for Vaccination-Only and Free Testing Models ($C_V$, $C_{V'}$, & $C_{V^\dagger}$) 6
- E. Models with Non-Free Testing ($C_{VT'}$, $C_{VT^\dagger}$) 7
- F. Multiple Testing ($C_{VLT'}$, $C_{VLT^\dagger}$) 8
- G. Comparison to Vaccination Without Testing 11

III. Exposure Model 13
- A. Alternative Exposure Models 13
- B. Fitting Force of Exposure Model 14
- C. Serosurvey Fits 16
- D. Context Sensitivities 16

IV. Context Economics 17

References 19


### LIST OF TABLES


I Data from L'Azou (2016). 16

II Data from Morrison (2010). 16

III Fits to seroprevalence data. 17

IV Sensitivity categories, $\mathbf{S}^{+}, \rho_H, \log_{10}\mathrm{OR}$. 17


### LIST OF FIGURES


1 **Outcomes for Status Quo**. Pathways show the probability of various life trajectories and ultimate costs for Eq. 1. 5

2 **Outcomes for Unconditional Vaccination**. Pathways show the probability of various life trajectories and ultimate costs for Eq. 2. Compared to Fig. 1, individuals that will experience infections over life are now either vaccinated before or after their first infections, meaning they may have a first-like infection converted to a second-like infection, or possibly avoid second-like infections. The vaccination may also be wasted, if given to individuals having already experienced multiple infections. 5

3 **Outcomes for Vaccination with a Free Ordinal Test**. Pathways show the probability of various life trajectories and ultimate costs for Eq. 3. Compared to Fig. 2, we never induced second-like infections, and indeed avoid all of them that would have occurred after the routine testing age. 6

4 **Outcomes for Vaccination with a Free Binary Test**. Pathways show the probability of various life trajectories and ultimate costs for Eq. 4. Note that the only distinction from Fig. 3 is the additional vaccination along the uppermost path, the trajectory corresponding to two or more infections prior to the routine vaccination age. While only a single term, this can make vaccination substantially more expensive in settings where multiple infections are frequent prior to the routine testing age. 7

5 **Maximum Vaccine Cost Fraction Allowing Net Benefit**. The approximate maxima for $\nu$ based on Eq. 8-9, including their convergence at younger ages. These curves represent the thresholds for vaccine costs given *real* tests, *i.e.* non-free with less than perfect sensitivity and specificity. Note that these curves only represent the restriction on $\nu$ for *some* benefit; the amount of benefit is generally shrinking faster than the cap on vaccine cost, particularly for the ordinal test. We facet by epidemiological parameters here: seroprevalence in 9-year-olds (a surrogate for dengue transmission level) and disparity (a measure of risk heterogeneity). . . . . 8
6 **Vaccine and Single Test Cost Fraction Boundary**. The approximate boundary for $\nu$ and $\tau$ based on Eq. 15-16. To be net beneficial, the costs of testing and vaccine (under a single test strategy) must result in a point below the relevant line (based on routine vaccination age and context). We show how the threshold line shifts with age. We facet by environmental sensitivity parameters here: seroprevalence in 9-year-olds (a surrogate for dengue transmission level) and disparity (a measure of risk heterogeneity). . . . . 9
7 **Outcomes for Vaccination with Multiple Ordinal Tests**. Pathways show the probability of various life trajectories and ultimate costs for Eq. 17. Unlike to single testing, both the upper (corresponding to infection(s) prior to testing) and lower (an initial non-passing test followed by any future trajectory) paths are allowed. . . . . 9
8 **Outcomes for Vaccination with Multiple Binary Tests**. Pathways show the probability of various life trajectories and ultimate costs for Eq. 18. Again, note that the only distinction from Fig. 7 is the additional vaccination along the uppermost path, the trajectory corresponding to two or more infections prior to the routine vaccination age. . . . . 10
9 **Vaccination Benefit with and without Testing, F/S=0.5**. . . . . 12
10 **Vaccination Benefit with and without Testing, F/S=0.2**. . . . . 12
11 **Trends by Age of Relevant Population States**. Assorted ${}_N\mathcal{P}_X$ trend lines, faceted by seroprevalence in 9-year-olds (a surrogate for overall force of exposure) and disparity level (a representation of population heterogeneity in exposure risk). . . . . 14

### I. ASSUMPTIONS

Based on our understanding of how CYD-TDV works in transmission settings of interest, we model the expected life time benefits and costs of vaccination interventions versus status quo on an individual basis. This framing implies that **interventions have no effect beyond the individual**, which effectively ignores any changes to transmission. Similarly, all **benefits and cost are on the margin**, so for example expenditure on testing and vaccination is independent of coverage level. However, economies of scale could be incorporated by, say, adjusting the vaccine or test prices to account for lower unit costs.

Our model is on a **discrete, annual time scale**. Practically, this means representing all dengue natural history effects as discrete-year effects (*e.g.* transient cross-serotype immunity). Likewise, interventions only occur once a year, and all data is interpreted at this yearly scale.

We assume **dengue is an environmental risk**, rather than a dynamically spread infection, with multiple, but indistinguishable, serotypes. Combined with the annual time scale, this means we represent dengue exposure risk as a probability per year-of-life effect; see SI Section *Exposure Model* for details. Since we assume the **dengue serotypes are indistinguishable**, they have identical infection probability, disease risk, circulation probability, *etc.* We assume that any particular serotype is life-time immunizing against that serotype, and that infection in one year precludes infection by any serotype in the next year. Finally, we assume no direct age-dependence for any aspect of dengue disease.

Using this model of dengue exposure, we pre-compute sample life trajectories for 1 million individuals (to ensure relatively smooth estimates), all living to age 70. We did not explicitly explore sensitivity to this life expectancy (either mean or distribution), but the qualitative trends can be determined from reasoning: increase in life expectancy increases the intervention value (and *vice versa*), as more people are likely to have multiple life infections, and any increase in single infections is well outside the test and vaccinate window. However, this effect applies to such a small proportion—those with a single infection by or during the test-then-vaccinate window, on the margin of having one or two total life time infections that gain or lose an infection by extending or shortening life decades later—and so we expect it to have little impact on any of these assessments.

For dengue disease natural history, we assume only the first and second infections have potentially associated disease.

We assume that, aside from infection-history-derived disease outcomes, vaccination has no impact on infections over life. That is, the transient immunity observed during CYD-TDV trials does not change the number of lifetime infections any individual experiences. We justify this based on the relatively short duration of immunity compared to life expectancy. By making this assumption, we can use pre-sampled life infection trajectories without having to perturb them by vaccination.

Finally, we assume tests are perfectly sensitive and specific. This provides an upper bound on benefit, which can be used to preclude combinations of strategy and price for vaccines and tests, irrespective of test performance. The detailed cost performance of real tests will vary by setting, test properties, and intervention approach. We assume any extant individual test history (*e.g.* previous lab confirmed dengue infection) is irrelevant to this strategy on a longer term basis, and so we ignore that testing; going forward, any non-vaccine related testing could be reasonably considered part of our cost estimate.

On balance, these assumptions tend to undervalue vaccination, while overvaluing testing. We judge that the net effect is likely small relative to estimating limiting space of a test-then-vaccinate regimen for the yes or no decision many settings now face for CYD-TDV. For those settings where the prices are in (or close enough to) the beneficial space, this type of analysis should be revisited with more modelling detail.

### II. COST-BENEFIT EQUATIONS

#### A. Definitions

Our model represents the following costs:

- $F$, $S$: the average individual costs of first-like and second-like infections, respectively. These should be based on local economic data, *e.g.* hospital costs, lost productivity, willingness to pay, but like most economic decisions will have a subjective element

- $V$, $T$: the unit cost of vaccine and testing regimens; these values are to be constrained, or have proposed prices to be evaluated against those constraints

- $\nu = \frac{V}{S}$, $\tau = \frac{T}{S}$: the costs of the vaccine and test as a fraction of the cost of second-like infection; deriving equations in these terms allows general results
- $C_X$, the total individual cost for scenario $X$ (no intervention, vaccination without testing, *etc.*)
- $\Delta_X$, the difference between status quo ($C_0$; no vaccination) and intervention $X$: $\Delta_X = C_0 - C_X$. For an intervention to have net benefit, $\Delta_X \geq 0$ must be true

Note, that these costs may represent any perspective (individual, societal, *etc.*), but should be from a consistent perspective.

The population is stratified by total lifetime infections, as well as number of infections at the routine testing age (or vaccination age, for the comparison scenario without testing). These proportions in turn determine the costs and benefits of a particular vaccination and testing strategy. We represent these proportions with the following general constructions:

- $\mathcal{P}_X$: proportion having $X$ infections during life; this value is age independent
- ${}_N\mathcal{P}_X\{A\}$, proportion having $N$ infections prior to consideration for vaccination at age $A$, and $X$ total lifetime infections
- $N$ and $X$ can include modifiers, like $X^+$ to mean $X$ or more, or have $\forall$ to mean any number of total infections (by definition, $X \geq N$).
- $\mathcal{C}\{A\}$, the conditional probability of seroconverting between age $A$ and $A+1$, given seronegativity at age $A$.

In general, we drop $A$, as it is the same for all terms in most equations. We only use it explicitly for multiple testing scenarios. Here are some example proportions which are relevant to our final derivation results:

The proportion of individuals that

- $\mathcal{P}_0$: ... are lifetime seronegative
- $\mathcal{P}_{1^+}$: ... have one or more lifetime infections
- ${}_0\mathcal{P}_{2^+}$, ${}_1\mathcal{P}_{2^+}$: ... have no or one infection, respectively, prior to routine age, and have two or more infections over life
- ${}_{2^+}\mathcal{P}_\forall$: ... have two or more infections at the time of vaccination
- ${}_1\mathcal{P}_\forall$, ${}_{1^+}\mathcal{P}_\forall$: ... have exactly one or at least one infection at the time of vaccination

In a later section, we will define an exposure model to estimate these probabilities, but we may take the specifics of that for granted while we layout the cost relationships. Because the cost model is defined only in terms of the probabilities, we could replace the exposure model with another approach that provides the same probabilities, as long as it addressed other pertinent assumptions, like the independence of individuals.

#### B. Status Quo and Vaccination Only Costs ($C_0$, $C_V$)

Without vaccination, an individual's expected lifetime dengue burden would be:

$$C_0 = (1 - \mathcal{P}_0)F + \mathcal{P}_{2^+}S \tag{1}$$

Note that the proportions are all independent of age, and only concern lifetime outcomes. When considering interventions, age becomes a factor (implicit with the ${}_X\mathcal{P}_N$ terms, which are a function of $A$). With universal vaccination (*i.e.*, irrespective of serostatus), the cost would be:

$$C_V = {}_1\mathcal{P}_\forall F + {}_{2^+}\mathcal{P}_\forall(F + S) + V + {}_0\mathcal{P}_{1^+}S \tag{2}$$

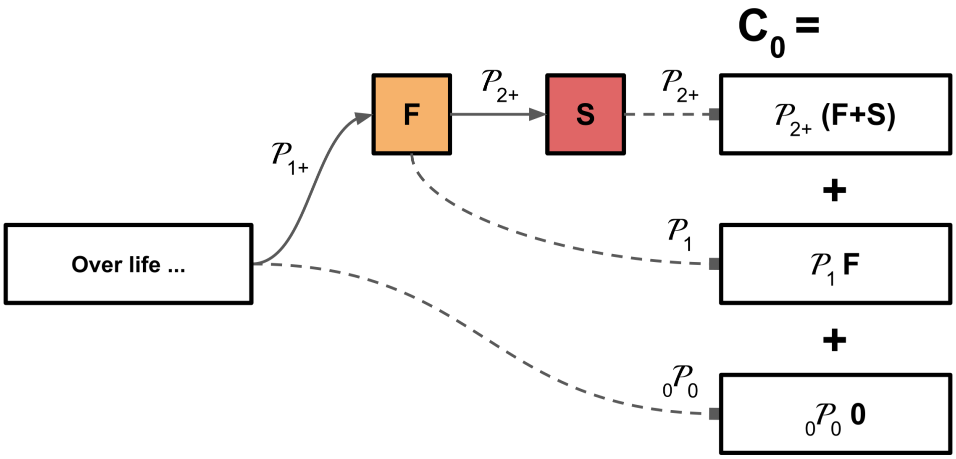


FIG. 1. **Outcomes for Status Quo**. Pathways show the probability of various life trajectories and ultimate costs for Eq. 1.

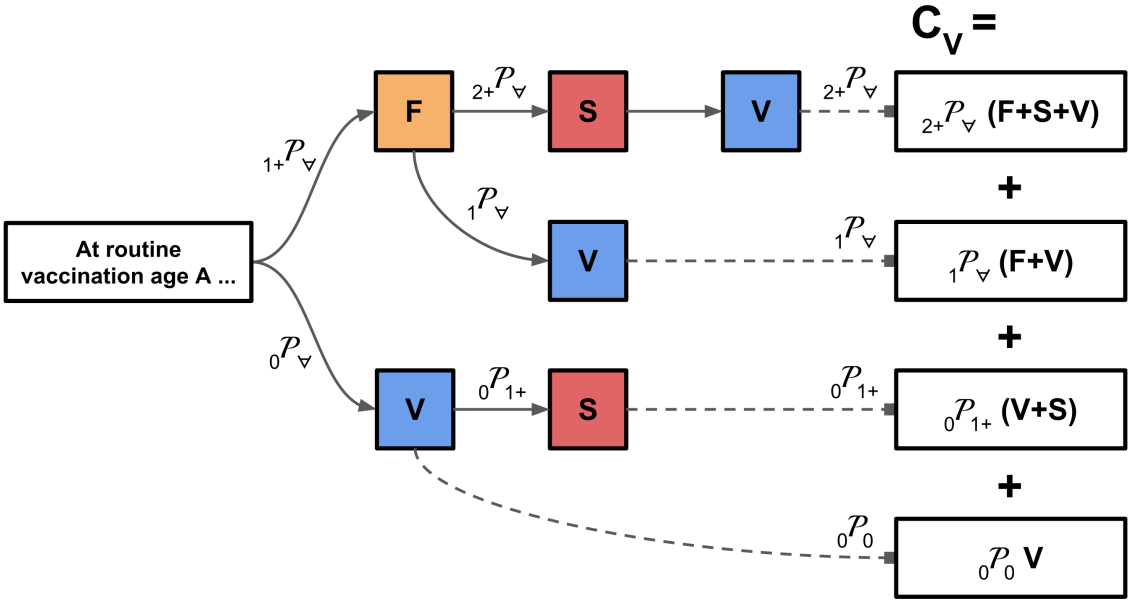


FIG. 2. **Outcomes for Unconditional Vaccination**. Pathways show the probability of various life trajectories and ultimate costs for Eq. 2. Compared to Fig. 1, individuals that will experience infections over life are now either vaccinated before or after their first infections, meaning they may have a first-like infection converted to a second-like infection, or possibly avoid second-like infections. The vaccination may also be wasted, if given to individuals having already experienced multiple infections.

#### C. Model with Free Testing

To begin understanding the cost model with testing, we imagine combining the vaccine with a free test (denoted $V'$); this assumption is obviously ludicrous—the test will cost resources to produce and use—but provides insight about the limits of vaccine value for an epidemiological setting.

We assume this imaginary test identifies the *number* of past dengue infections, so we refer to it as the *ordinal* test. This test enables us to only vaccinate seropositives that *could* have another disease-bearing infection, and avoid vaccinating seronegatives until they seroconvert. We use the test every year from the routine testing age, until the individual receives the vaccine. The cost for this hypothetical regimen is:

$$C_{V'} = (1 - \mathcal{P}_0)F + (1 - \mathcal{P}_0 - {}_{2+}\mathcal{P}_\forall)V' + {}_{2+}\mathcal{P}_\forall S \tag{3}$$

Note that $V'$ vaccination only occurs for seropositives that have only had one infection and individuals that se-

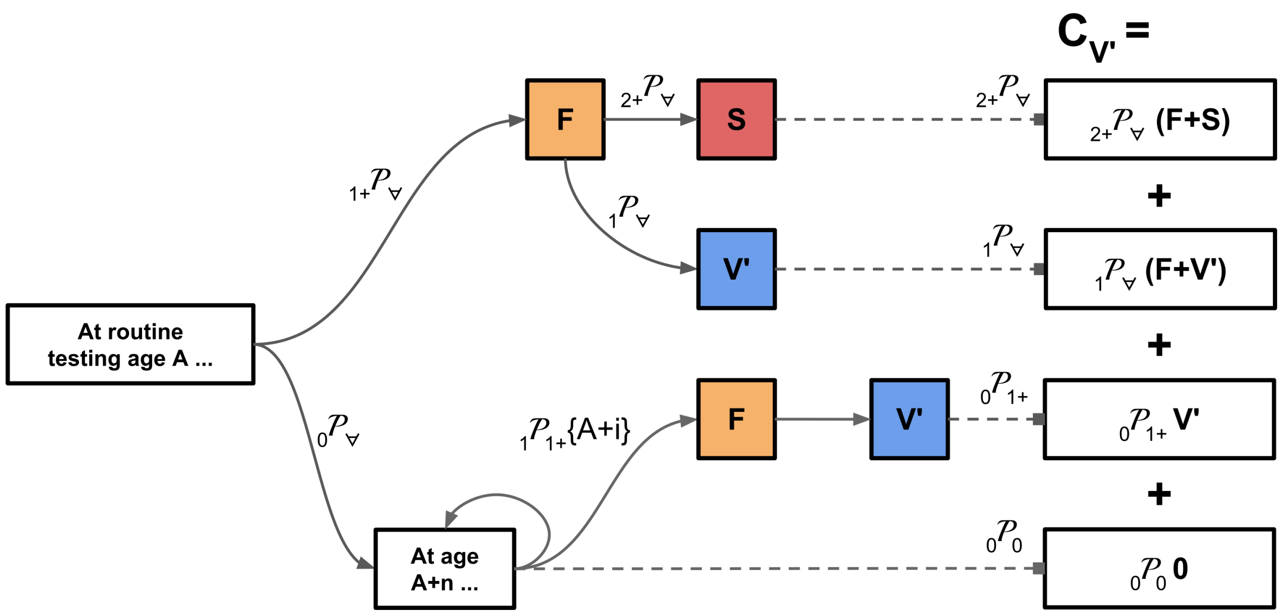


FIG. 3. **Outcomes for Vaccination with a Free Ordinal Test**. Pathways show the probability of various life trajectories and ultimate costs for Eq. 3. Compared to Fig. 2, we never induced second-like infections, and indeed avoid all of them that would have occurred after the routine testing age.

roconvert after the routine testing age; because we assume no benefit against third and later infections, we do not administer vaccine to those with multiple past infections. If we have a test that only detects seropositivity but not detailed infection history, which we refer to as a *binary* test, then costs would be:

$$C_{V^\dagger} = (1-\mathcal{P}_0)F + (1-\mathcal{P}_0)V^\dagger + {}_{2^+}\mathcal{P}_\forall S \tag{4}$$

which we distinguish by the $V^\dagger$ term. The only change is on the coefficient for the $V$ term (and the $\nu = \frac{V}{S}$ terms, when we switch to non-dimensional form); this result will be consistent as we consider more complicated models.

#### D. Cost-Benefit Constraint Equations for Vaccination-Only and Free Testing Models ($C_V$, $C_{V'}$, & $C_{V^\dagger}$)

If we compare all these testing scenarios to non-vaccination:

$$\begin{aligned}
\Delta_V = C_0 - C_V &= (1-\mathcal{P}_0 - {}_1\mathcal{P}_\forall - {}_{2^+}\mathcal{P}_\forall)F + (\mathcal{P}_{2^+} - {}_0\mathcal{P}_{1^+} - {}_{2^+}\mathcal{P}_\forall)S - V \\
&= (\mathcal{P}_{1^+} - {}_1\mathcal{P}_\forall - {}_{2^+}\mathcal{P}_\forall)F + ({}_0\mathcal{P}_{2^+} - {}_0\mathcal{P}_{1^+} + {}_1\mathcal{P}_{2^+} + {}_{2^+}\mathcal{P}_{2^+} - {}_{2^+}\mathcal{P}_\forall)S - V \\
&= {}_0\mathcal{P}_{1^+}F + ({}_1\mathcal{P}_{2^+} - {}_0\mathcal{P}_1)S - V \qquad (5)\\
\Delta_{V'} &= (\mathcal{P}_{2^+} - {}_{2^+}\mathcal{P}_\forall)S - (1-\mathcal{P}_0 - {}_{2^+}\mathcal{P}_\forall)V' \qquad (6)\\
\Delta_{V^\dagger} &= (\mathcal{P}_{2^+} - {}_{2^+}\mathcal{P}_\forall)S - (1-\mathcal{P}_0)V^\dagger \qquad (7)
\end{aligned}$$

Recall that by definition, vaccination without testing is only net beneficial if $\Delta_V \geq 0$. Testing can expand the region of vaccine benefit, but only if $\Delta_{V'} \geq 0$ or $\Delta_{V^\dagger} \geq 0$, depending on the test mechanism. For routine vaccination at a certain age to be net beneficial after the introduction of any real testing cost, the vaccine regimen cost must obey:

$$\nu' < \frac{\mathcal{P}_{2^+} - {}_{2^+}\mathcal{P}_\forall}{1-\mathcal{P}_0 - {}_{2^+}\mathcal{P}_\forall} \tag{8}$$

$$\nu^\dagger < \frac{\mathcal{P}_{2^+} - {}_{2^+}\mathcal{P}_\forall}{1-\mathcal{P}_0} \tag{9}$$

Note that combined with either of the free tests, the constraint on vaccine cost depends only on three generic parameters of the context (the average cost of second-like infections ($S$, implicit within $\nu$), and the lifetime probabilities

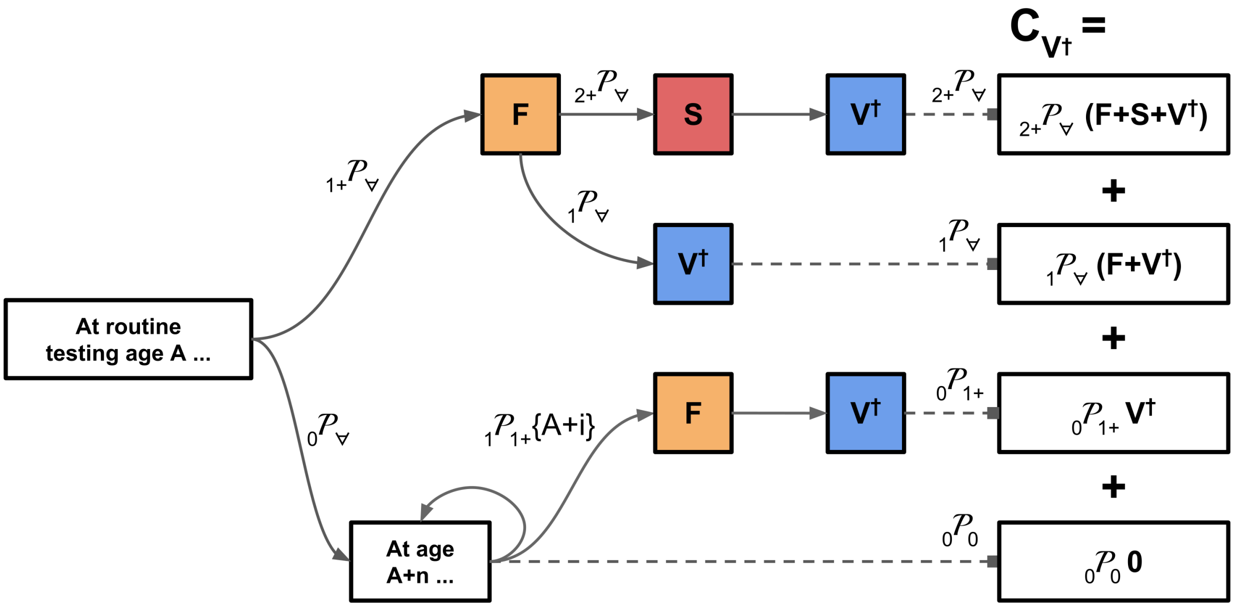


FIG. 4. **Outcomes for Vaccination with a Free Binary Test**. Pathways show the probability of various life trajectories and ultimate costs for Eq. 4. Note that the only distinction from Fig. 3 is the additional vaccination along the uppermost path, the trajectory corresponding to two or more infections prior to the routine vaccination age. While only a single term, this can make vaccination substantially more expensive in settings where multiple infections are frequent prior to the routine testing age.

of 0 or $2^+$ infections) and one parameter of the intervention (*i.e.*, age $A$ of consideration for vaccination, which determines the fraction of the population that has had two or more infections, $_{2^+}\mathcal{P}_\forall$, at that point). As the routine age is shifted earlier, the probability of two infections prior to consideration for vaccination becomes vanishingly small, *i.e.* $_{2^+}\mathcal{P}_\forall \to 0$. Therefore, at a young enough vaccine age the constraint equations converge to the ratio

$$\nu' = \nu^\dagger \leq \frac{\mathcal{P}_{2^+}}{1 - \mathcal{P}_0} \tag{10}$$

or the fraction of infectees that will have multiple infections out of all infectees. We could have arrived at this result reasoning from event probabilities: an individual only pays the vaccine cost if they experience at least one infections $(1 - \mathcal{P}_0)$, and only individuals that experience multiple infections benefit (by avoiding $S$). Therefore, the maximum value of $V$ is the conditional probability of experiencing $S$ given any infections.

Note that $\nu' \geq \nu^\dagger$ for any routine testing age, but the value of $\nu^\dagger$ grows faster with decreasing age (as it must for them to converge).

#### E. Models with Non-Free Testing ($C_{VT'}$, $C_{VT^\dagger}$)

If we now acknowledge the cost of testing, $T$, and consider only a single test (instead of potentially testing several times), the individual costs are:

$$C_{VT'} = T' + (1 - \mathcal{P}_0)F + ({_0}\mathcal{P}_{2^+} + {_{2^+}}\mathcal{P}_\forall)S + {_1}\mathcal{P}_\forall V \tag{11}$$
$$C_{VT^\dagger} = T^\dagger + (1 - \mathcal{P}_0)F + ({_0}\mathcal{P}_{2^+} + {_{2^+}}\mathcal{P}_\forall)S + {_{1^+}}\mathcal{P}_\forall V \tag{12}$$

and the intervention benefits are:

$$\begin{aligned}\Delta_{VT'} &= (\mathcal{P}_{2^+} - {_0}\mathcal{P}_{2^+} - {_{2^+}}\mathcal{P}_\forall)S - T' - {_1}\mathcal{P}_\forall V \\ &= \mathcal{P}_{2^+}{_1}\mathcal{P}_{2^+}S - T' - {_1}\mathcal{P}_\forall V \end{aligned} \tag{13}$$
$$\Delta_{VT^\dagger} = \mathcal{P}_{2^+}{_1}\mathcal{P}_{2^+}S - T^\dagger - {_{1^+}}\mathcal{P}_\forall V \tag{14}$$

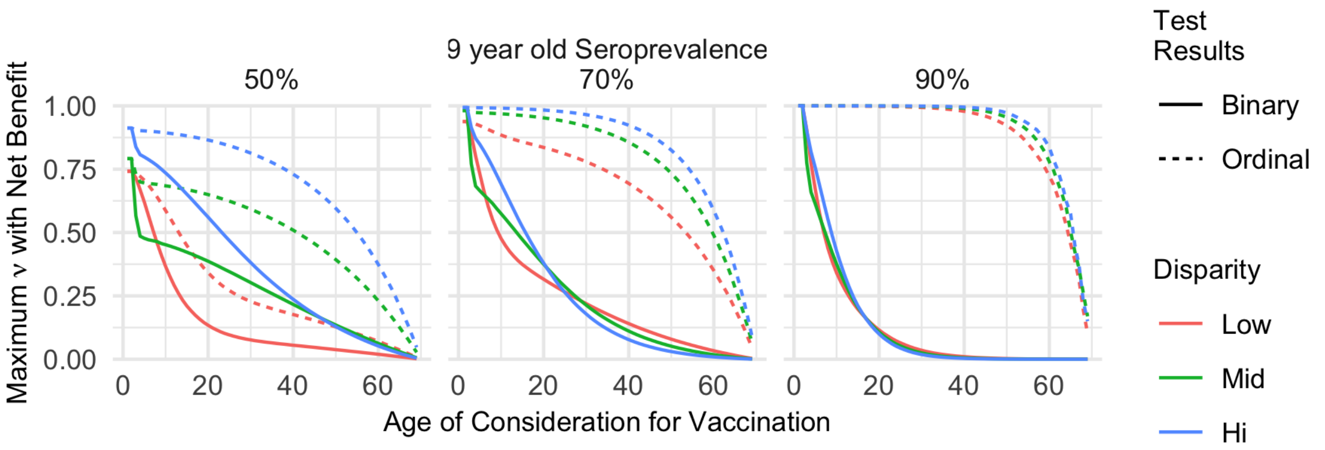


FIG. 5. **Maximum Vaccine Cost Fraction Allowing Net Benefit**. The approximate maxima for $\nu$ based on Eq. 8-9, including their convergence at younger ages. These curves represent the thresholds for vaccine costs given *real* tests, *i.e.* non-free with less than perfect sensitivity and specificity. Note that these curves only represent the restriction on $\nu$ for *some* benefit; the amount of benefit is generally shrinking faster than the cap on vaccine cost, particularly for the ordinal test. We facet by epidemiological parameters here: seroprevalence in 9-year-olds (a surrogate for dengue transmission level) and disparity (a measure of risk heterogeneity).

Which makes our non-free, single testing constraints for cost effectiveness:

$$\tau' + {}_1\mathcal{P}_{\forall}\nu \leq \mathcal{P}_{2^+1}\mathcal{P}_{2^+} \tag{15}$$

$$\tau^{\dagger} + {}_{1^+}\mathcal{P}_{\forall}\nu \leq \mathcal{P}_{2^+1}\mathcal{P}_{2^+} \tag{16}$$

Notably, there is still no dependence on the cost of first infections ($F$).

#### F. Multiple Testing ($C_{VLT'}$, $C_{VLT^{\dagger}}$)

Now consider if we allow testing every year, up to a maximum of $L$ tests (including the first). Like the free test intervention, we may identify people who seroconvert after initial consideration for vaccination, though since there is a test cost, there will be diminishing returns (and eventually losses) with repeat testing.

Relative to $C_{VT'}$ or $C_{VT^{\dagger}}$, lifetime seronegatives ($\mathcal{P}_0$) will pay for $L-1$ additional tests. The fraction of the population in ${}_0\mathcal{P}_{1^+}$ will pay for some number of additional tests, and possibly vaccination (*i.e.*, for the proportion of individuals seroconverting during the testing window). This means costs for this strategy will change some ${}_0\mathcal{P}_{1^+}\{A\}$ (*i.e.*, seronegative at $A$, the routine initial testing age) to ${}_1\mathcal{P}_{1^+}\{A+n\}$ at some later age prior to $A+L-1$ (*i.e.*, seroconverting during the testing window, and therefore vaccinated).

Relative to single testing, this will add some average number of tests and expand vaccination based on the amount of seroconversion during the testing period, thus increasing the cost of the intervention. In return, there will be a reduction in the proportion incurring $S$. Relative to indefinite free testing, this reduction will be incomplete, as some proportion will pass the entire $L$ years of the testing period without seroconversion, but still suffer multiple future infections. Isolating just this difference relative to single tests, we define a substitute $f$ which represents all the costs with respect to $L$:

$$C_{VLT'} = C_{VT'} + \left({}_0\mathcal{P}_{2^+}\{A+L-1\} - {}_0\mathcal{P}_{2^+}\{A\}\right)S + f(L) \tag{17}$$

$$C_{VLT^{\dagger}} = C_{VT^{\dagger}} + \left({}_0\mathcal{P}_{2^+}\{A+L-1\} - {}_0\mathcal{P}_{2^+}\{A\}\right)S + f(L) \tag{18}$$

The $S$ burden is reduced from the seronegative population (with multiple future infections) when first considered for vaccination, to the fraction of that group that remains un-infected through the whole testing period. In exchange, we add some new vaccination and testing costs, $f(L)$. Note that $f$ is structurally identical for both binary and ordinal tests; since all tests after the first are only applied to the proportion that was seronegative in the previous year, and since the model permits at most one infection per year, the tests are only on seronegative or first-infection

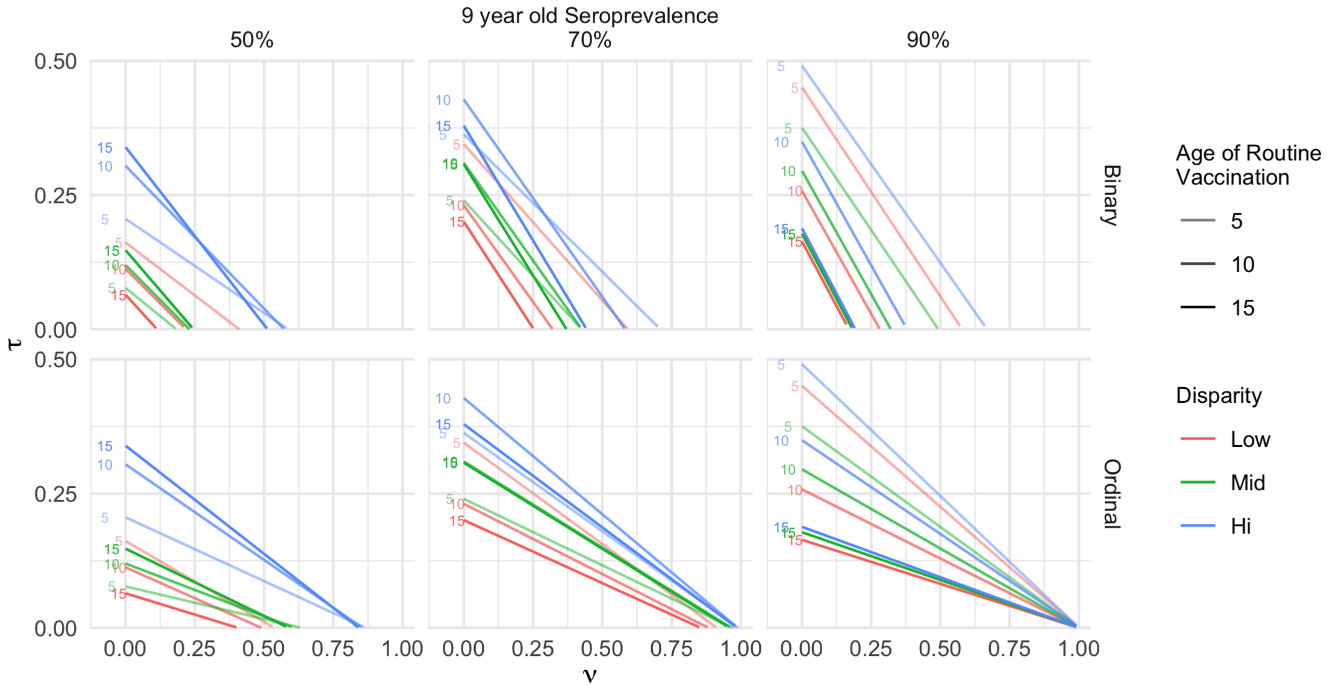


FIG. 6. **Vaccine and Single Test Cost Fraction Boundary**. The approximate boundary for $\nu$ and $\tau$ based on Eq. 15-16. To be net beneficial, the costs of testing and vaccine (under a single test strategy) must result in a point below the relevant line (based on routine vaccination age and context). We show how the threshold line shifts with age. We facet by environmental sensitivity parameters here: seroprevalence in 9-year-olds (a surrogate for dengue transmission level) and disparity (a measure of risk heterogeneity).

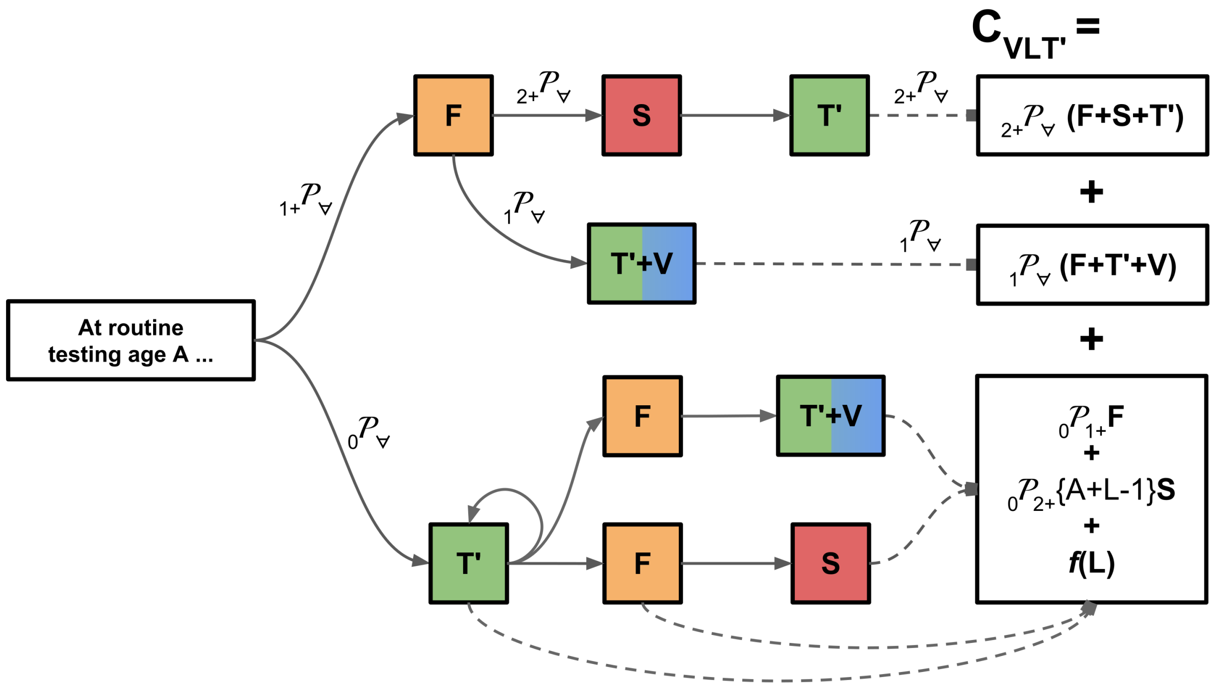


FIG. 7. **Outcomes for Vaccination with Multiple Ordinal Tests**. Pathways show the probability of various life trajectories and ultimate costs for Eq. 17. Unlike to single testing, both the upper (corresponding to infection(s) prior to testing) and lower (an initial non-passing test followed by any future trajectory) paths are allowed.

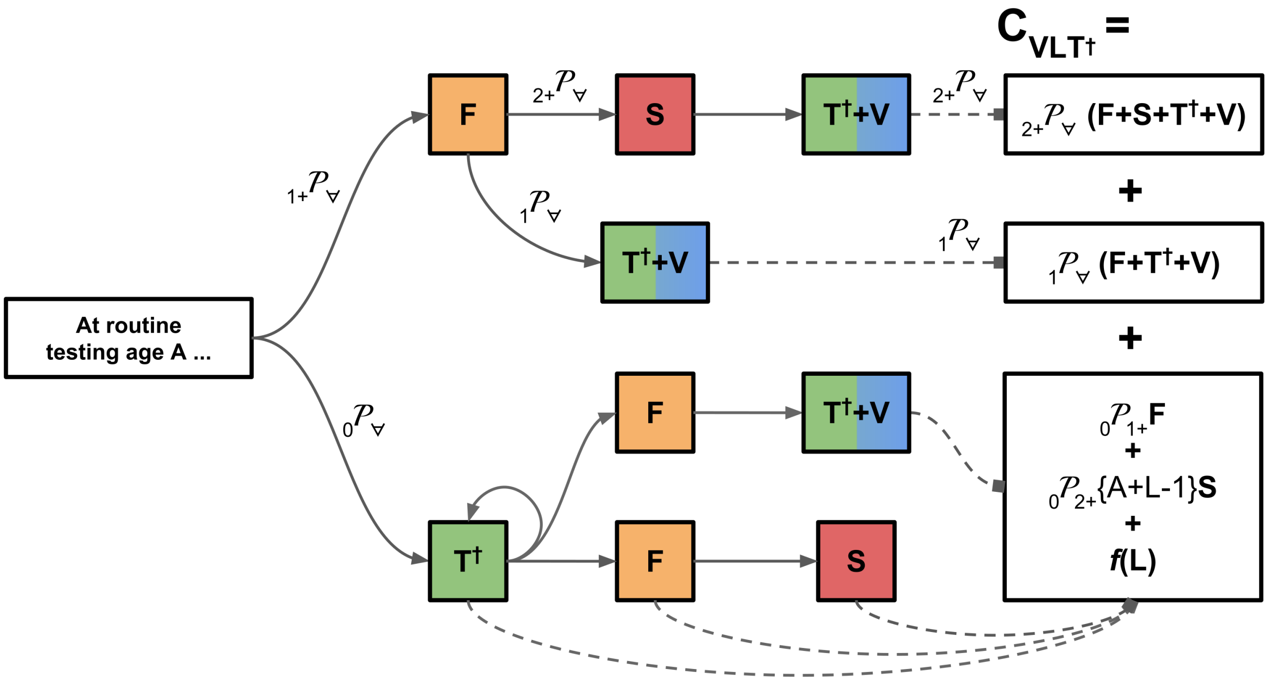


FIG. 8. **Outcomes for Vaccination with Multiple Binary Tests**. Pathways show the probability of various life trajectories and ultimate costs for Eq. 18. Again, note that the only distinction from Fig. 7 is the additional vaccination along the uppermost path, the trajectory corresponding to two or more infections prior to the routine vaccination age.

seropositives. Therefore the test action is indistinguishable for all individuals re-testing, though this is only true assuming the test provides perfect information.

We define $f$ as:

$$
\begin{aligned}
f(L) = &\left(\mathcal{P}_0 + {}_0\mathcal{P}_{1^+}\{A{+}L{-}1\}\right)(L{-}1)T + {}_0\mathcal{P}_{1^+}\{A\}\times \\
&\Bigg(\mathcal{C}\{A\}(V{+}T) + \mathcal{C}\{A{+}1\}\frac{{}_0\mathcal{P}_{1^+}\{A{+}1\}}{{}_0\mathcal{P}_{1^+}\{A\}}(V{+}2T) + \mathcal{C}\{A{+}2\}\frac{{}_0\mathcal{P}_{1^+}\{A{+}2\}}{{}_0\mathcal{P}_{1^+}\{A\}}(V{+}3T) + \dots \\
&\quad \mathcal{C}\{A{+}L{-}2\}\frac{{}_0\mathcal{P}_{1^+}\{A{+}L{-}2\}}{{}_0\mathcal{P}_{1^+}\{A\}}(V{+}(L{-}1)T)\Bigg)
\end{aligned}
\tag{19}
$$

which can be conveniently expressed in terms of a summation:

$$
f(L) = {}_0\mathcal{P}_{\forall}\{A{+}L{-}1\}(L{-}1)T + \sum_{i=0}^{L-2}\mathcal{C}\{A{+}i\}{}_0\mathcal{P}_{1^+}\{A{+}i\}\left(V{+}(i{+}1)T\right)
\tag{20}
$$

Note that $f(L{=}1) = 0$, since the summation term is empty. This definition of $f(L)$ can then be used in the net benefit relationships:

$$
\begin{aligned}
\Delta_{VLT'} &= \Delta_{VT'} - \left({}_0\mathcal{P}_{2^+}\{A+L-1\} - {}_0\mathcal{P}_{2^+}\{A\}\right)S - f(L) \\
&= \left(\mathcal{P}_{2^+} - {}_{2^+}\mathcal{P}_{\forall} - {}_0\mathcal{P}_{2^+}\{A+L-1\}\right)S - T' - {}_1\mathcal{P}_{\forall}V - f(L)
\end{aligned}
\tag{21}
$$

$$
\Delta_{VLT^\dagger} = \left(\mathcal{P}_{2^+} - {}_{2^+}\mathcal{P}_{\forall} - {}_0\mathcal{P}_{2^+}\{A+L-1\}\right)S - T^\dagger - {}_{1^+}\mathcal{P}_{\forall}V - f(L)
\tag{22}
$$

Again, the cost of first-like infections, $F$, does not appear anywhere in the constraints. We can also divide by $S$ and again work in the non-dimensional terms, with $\hat{f}$ the non-dimensional version of $f$:

$$\hat{f}(L) = {}_0\mathcal{P}_\forall\{A{+}L{-}1\}(L{-}1)\tau + \sum_{i=0}^{L-2} \mathcal{C}\{A{+}i\}_0\mathcal{P}_{1^+}\{A{+}i\}\left(\nu{+}(i{+}1)\tau\right) \tag{23}$$

$$\tau' + {}_1\mathcal{P}_\forall\nu + \hat{f}(L) \leq \left(\mathcal{P}_{2^+} - {}_{2^+}\mathcal{P}_\forall - {}_0\mathcal{P}_{2^+}\{A + L - 1\}\right) \tag{24}$$

$$\tau^\dagger + {}_{1^+}\mathcal{P}_\forall\nu + \hat{f}(L) \leq \left(\mathcal{P}_{2^+} - {}_{2^+}\mathcal{P}_\forall - {}_0\mathcal{P}_{2^+}\{A + L - 1\}\right) \tag{25}$$

#### G. Comparison to Vaccination Without Testing

As a final step, we compare the benefits of vaccination with and without testing. This allows us to determine where adding testing of makes vaccination a more cost-effective intervention. Recalling $C_V$ and $C_{VLT^\dagger}$ from Eq. 2 and Eq. 18, respectively, we can write:

$$C_V - C_{VLT^\dagger} = -{}_0\mathcal{P}_{1^+}F + (1 - {}_{1^+}\mathcal{P}_\forall)V + \left({}_0\mathcal{P}_{1^+} - {}_0\mathcal{P}_{2^+}\{A + L - 1\}\right)S - T^\dagger - f(L) \tag{26}$$

For testing to improve the intervention cost-benefit, $C_V - C_{VLT^\dagger} \geq 0$. Imposing that constraint, and re-arranging to match the form of Eq. 25:

$$\tau^\dagger + {}_{1^+}\mathcal{P}_\forall\nu + \hat{f}(L) \leq \left({}_0\mathcal{P}_{1^+} - {}_0\mathcal{P}_{2^+}\{A + L - 1\}\right) - {}_0\mathcal{P}_{1^+}\frac{F}{S} + \nu \tag{27}$$

Some notable results are derivable directly from this relationship. Increasing vaccine cost increases the advantage of the testing intervention, which is intuitive: testing decreases vaccination rates, so the more expensive the vaccine, the more cost avoided by testing. The other direct result is that increasing costs of first-like infections *decreases* the value of testing. This outcome can be understood by thinking about what the vaccine does in practice, namely preventing one of $F$ or $S$, and what testing does, namely shifting prevention from $F$ to $S$. As $F$ and $S$ become closer, the value to preventing $S$ instead of $F$ decreases, and thus so does value to testing. As explained in the **Context Economics** section, we consider a pessimistic $\frac{F}{S} = 0.5$.

In general, we see that including testing makes the intervention more beneficial in lower transmission settings, while it is wasted in high transmission settings. In high transmission settings, we already have a lot of information about serostatus, so testing provides little benefit.

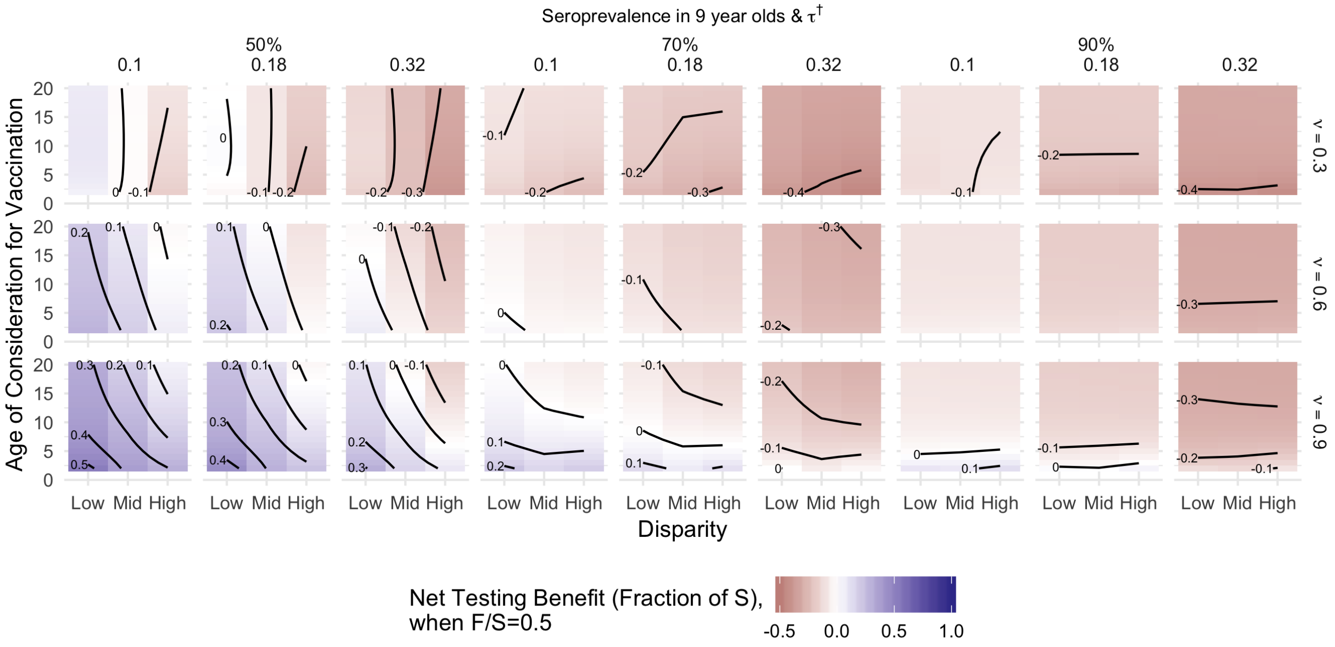


FIG. 9. **Vaccination Benefit with and without Testing, F/S=0.5**.

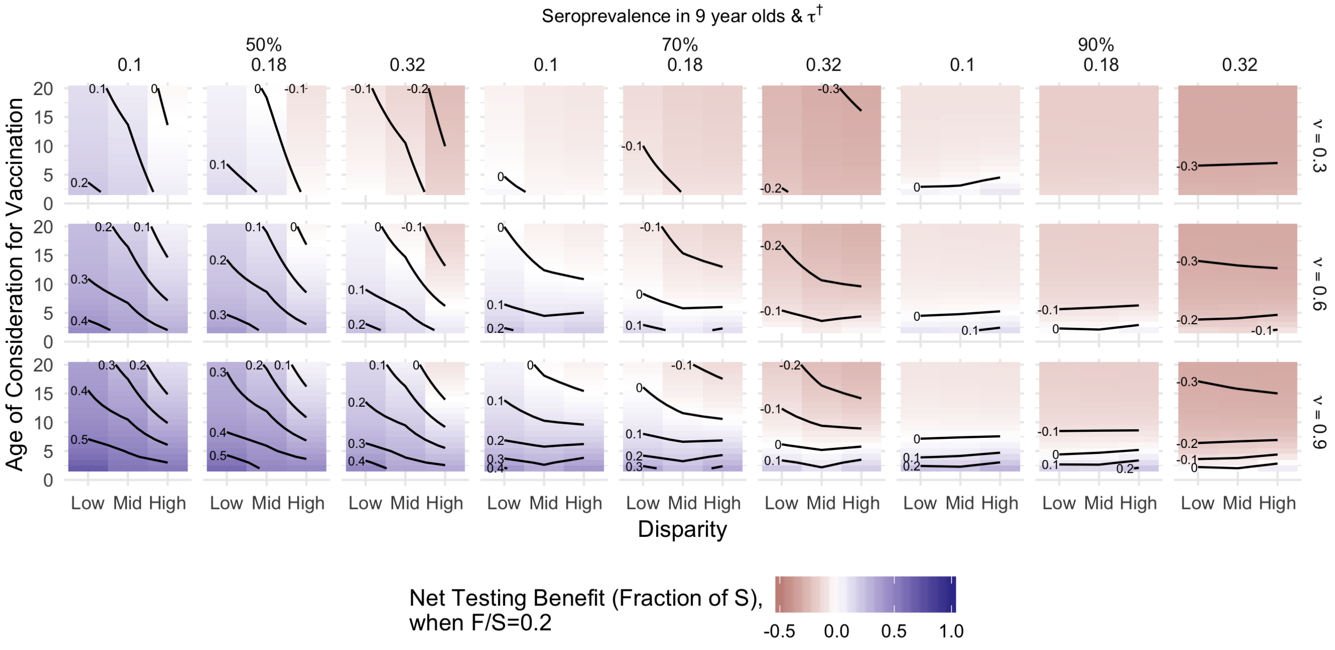


FIG. 10. **Vaccination Benefit with and without Testing, F/S=0.2**.

### III. EXPOSURE MODEL

As stated earlier, we assume there is a constant force-of-exposure. However, we also divide individuals into low or high risk sub-populations, so there are three fundamental parameters in the model: the two exposure risks and the proportion in high versus low risk.

For natural history of the infection, we assume that exposure to a serotype leads to infection if the individual has not been exposed to that serotype previously. We also assume that infection in one year precludes infection in the following year, consistent with empirical observations of temporary cross immunity. Finally, we assume only one dengue serotype is circulating in any given year. In the next section we consider relaxing these assumptions, and justify our decision to use these simpler assumptions.

We can use age-seroprevalence data to fit this force-of-exposure model. These data allow us to estimate the probability that individuals have experienced at least one infection by a given age. If we define the constant exposure forces (and complementary avoidance probabilities) $f_X = 1 - s_X$ and probability of being in the high risk group as $\rho_H$, the probability of being seropositive at age $A$ is:

$$\mathbf{P}(+|A) = \rho_H(1 - s_H^A) + (1 - \rho_H)(1 - s_L^A) \tag{28}$$

Once we fit that model to specific age-seroprevalence data, we can then use simulation to estimate the relevant ${}_N\mathcal{P}_X$ terms for that context, and thus the $\tau$-$\nu$ constraints for that setting. While analytical derivation of these is probably feasible for this set of assumptions, simulation affords us the ability to consider a broader range of assumptions in the future without having to re-derive outcomes.

Recall that for the particulars of our cost model, we need the following probabilities:

The proportion of individuals that

- ${}_0\mathcal{P}_\forall$: ... are seronegative at a given age
- $\mathcal{P}_{1^+}$: ... have one or more lifetime infections
- ${}_0\mathcal{P}_{2^+}$, ${}_1\mathcal{P}_{2^+}$: ... have no or one infection, respectively, prior to consideration for vaccination, and have two or more infections over life
- ${}_{2^+}\mathcal{P}_\forall$: ... have two or more infections at the time of vaccination
- ${}_1\mathcal{P}_\forall$, ${}_{1^+}\mathcal{P}_\forall$: ... have exactly one or at least one infection at the time of vaccination; and
- $\mathcal{C}\{A\}$, the conditional probability of seroconverting between age $A$ and $A + 1$, given seronegativity at age $A$.

#### A. Alternative Exposure Models

If there were more readily available data, it might be worth considering models incorporating (1) maternally-derived temporary immunity and (2) multiple serotype circulation.

To include maternal immunity, we would need to consider the probability of having a previously infected mother as part of the first exposure year. If an individual had an exposed mother, we would simply assume one the individuals was immune in that year, meaning one fewer exposure year in their life history. We could derive the probability of maternal exposure with maternal-age distribution and the age-seroprevalence from the fit, meaning we do not actually need an additional parameter, just more data.

For multiple simultaneously circulating serotypes, we considered allowing up-to four exposures per year, probabilistically ordered by relative weights. Since all the fitting is by first exposure, we could fit any one parameter model of exposure (*e.g.* a four-trial binomial draw). This would tend to drive up second infection rates, perhaps more accurately representing high risk populations in hyper-endemic locales where all four dengue serotypes routinely circulate. This would in turn tend to drive the benefit curve towards earlier consideration for routine vaccination. However, the data to correctly parametrise and constrain such a model (for example, age sero-ordinality surveys) is more detailed and does not appear to be commonly available.

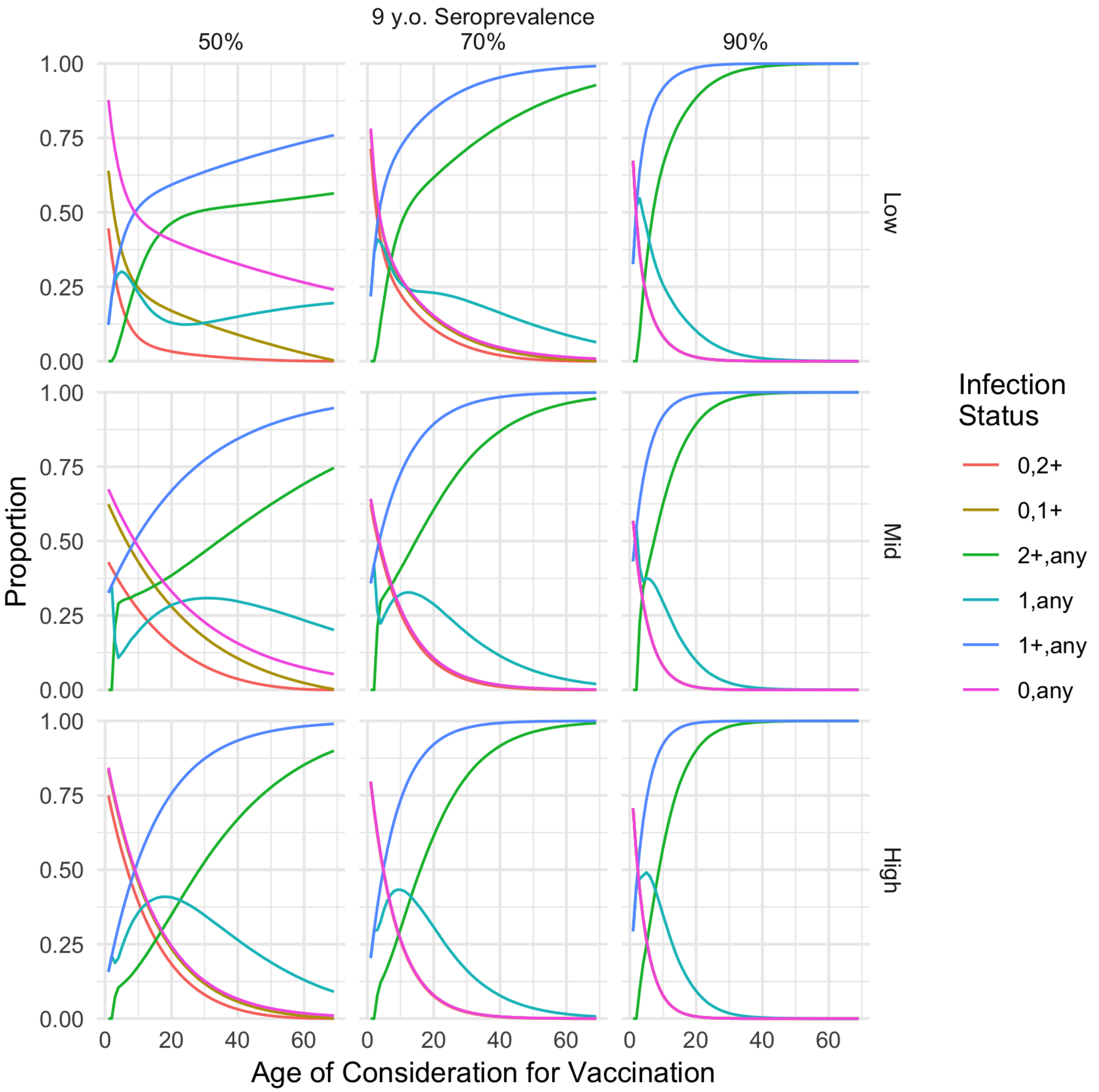


FIG. 11. **Trends by Age of Relevant Population States**. Assorted ${}_N\mathcal{P}_X$ trend lines, faceted by seroprevalence in 9-year-olds (a surrogate for overall force of exposure) and disparity level (a representation of population heterogeneity in exposure risk).

#### B. Fitting Force of Exposure Model

To fit the force of exposure model, we change parameters to break the parameter symmetry (*i.e.*, that either risk category could be higher or lower) and express them parameters so that they have infinite domain, which is preferable for most numerical solvers:

```
#' in R:
logit <- function(x) return(1/(1+exp(-x)))

#' @param A, the age being considered, scanned over fixed values
#' @param lgtfoeH, logit of high risk force of exposure
#' @param lgtRR, logit of the faorce of exposure relative risk for the low risk group
#' @param lgtpH, logit of the proportion of the population that is high risk
prob <- function(
  A, lgtfoeH, lgtRR, lgtpH
) {
  foeH <- logit(lgtfoeH)
  foeL <- logit(lgtRR)*foeH
  pH   <- logit(lgtpH)
  return(pH*(1-(1-foeH)^A) + (1-pH)*(1-(1-foeL)^A))
}
```

We can then use this function to calculate negative log-likelihoods for various parameter combinations, compared to the observed data, and optimize on those values.

#### C. Serosurvey Fits

Using this model, we fit to two data sets, Table I from the CYD14 trial [1] (reference Table 4; there is a discrepancy for Indonesia in 13-16 year-olds category when compared to reference Table 1; we have assumed Table 4 is correct) and Table II from work in Peru [2] (sample numbers read from reference Fig. 2 and percentiles extracted from reference Fig. 3 using a webtool [3]):

| Country | Age | N | $\mathbf{S}^+$ |
|---|---|---|---|
| Indonesia | 2-4 | 87 | 50 |
| | 5-8 | 90 | 76 |
| | 9-12 | 116 | 101 |
| | 13-16 | 57 | 56 |
| Malaysia | 2-4 | 68 | 22 |
| | 5-8 | 82 | 28 |
| | 9-12 | 102 | 59 |
| | 13-16 | 48 | 32 |
| Philippines | 2-4 | 150 | 87 |
| | 5-8 | 167 | 125 |
| | 9-12 | 157 | 139 |
| | 13-16 | 128 | 119 |
| Thailand | 2-4 | 68 | 33 |
| | 5-8 | 118 | 73 |
| | 9-12 | 117 | 93 |
| | 13-16 | 38 | 32 |
| Vietnam | 2-4 | 97 | 43 |
| | 5-8 | 142 | 70 |
| | 9-12 | 145 | 90 |
| | 13-16 | 22 | 17 |

TABLE I. Data from L'Azou (2016).

| Country | Age | N | $\mathbf{S}^+$ |
|---|---|---|---|
| Peru | 5 | 261 | 146 |
| | 6 | 307 | 209 |
| | 7 | 340 | 244 |
| | 8 | 368 | 269 |
| | 9 | 353 | 273 |
| | 10 | 340 | 273 |
| | 11 | 327 | 272 |
| | 12 | 320 | 257 |
| | 13 | 267 | 222 |
| | 14 | 238 | 209 |
| | 15 | 205 | 187 |
| | 16 | 147 | 135 |
| | 17 | 98 | 93 |

TABLE II. Data from Morrison (2010).

For our practical examples, we use Peru, a Latin American setting, and Malaysia, a Southeast Asian setting. The Peru data is ideal for the maximum-likelihood style approach we used. The aggregated CYD-14 data, on the other hand, throws away information and complicates model fitting, whether using maximum likelihood or other approaches. While the data may "look better" after aggregation, this mode of reporting in fact impedes further study.

#### D. Context Sensitivities

For our parameter sensitivities, we considered three dimensions: *high risk fraction*, *aggregate force of exposure*, and *disparity*. In keeping with past analyses, we distinguish aggregate force of exposure by seroprevalence at a particular age. We define disparity by the seroprevalence odds ratio (OR) between the high and low risk groups. These values are governed by:

$$
\begin{aligned}
\mathbf{S}^{+}(A) &= \rho_H \mathbf{S}^{+}_H(A) + (1-\rho_H)\mathbf{S}^{+}_L(A) \\
&= \rho_H(1-(1-f_H)^A) + (1-\rho_H)(1-(1-f_L)^A) \\
&= \rho_H(1-{s_H}^A) + (1-\rho_H)(1-{s_L}^A) \qquad (29)\\
\mathrm{OR} &= \frac{\mathbf{S}^{+}_H(A)(1-\mathbf{S}^{+}_L(A))}{\mathbf{S}^{+}_L(A)(1-\mathbf{S}^{+}_H(A))} = \frac{(1-{s_H}^A){s_L}^A}{(1-{s_L}^A){s_H}^A} \qquad (30)
\end{aligned}
$$

Therefore, for a particular $\{\rho_H, \mathbf{S}^{+} = 1-\mathbf{S}^{-}, \mathrm{OR}\}$, where the dependence on $A$ is implicit, we can identify $\alpha = {s_H}^A$ and $\beta = {s_L}^A$:

$$
\begin{aligned}
\alpha &= 1 - \frac{\mathbf{S}^{+}}{\rho_H} + \frac{1-\rho_H}{\rho_H} - \frac{1-\rho_H}{\rho_H}\beta \\
&= \frac{1-\mathbf{S}^{+}}{\rho_H} - \frac{1-\rho_H}{\rho_H}\beta = \frac{\mathbf{S}^{-}}{\rho_H} - \frac{1-\rho_H}{\rho_H}\beta \qquad (31)\\
\mathrm{OR} &= \frac{(1-\alpha)\beta}{(1-\beta)\alpha} \\
0 &= \mathrm{OR}\alpha + (1-\mathrm{OR})\alpha\beta - \beta \\
&= \mathrm{OR}(\mathbf{S}^{-} - (1-\rho_H)\beta) + (1-\mathrm{OR})(\mathbf{S}^{-} - (1-\rho_H)\beta)\beta - \rho_H\beta \\
&= \mathrm{OR}\mathbf{S}^{-} + ((1-\mathrm{OR})(\mathbf{S}^{-} - \rho_H) - \mathrm{OR})\beta - (1-\mathrm{OR})(1-\rho_H)\beta^2 \qquad (32)
\end{aligned}
$$

where Eq. 32 can be solved with the quadratic equation, subject to $\beta \in (0,1)$, and substituted into Eq. 31.

We are being somewhat circular with the concept of disparity, in that we fit the serosurvey data assuming a two-group risk model, and then calculated the OR, rather than identifying mechanistically a useful distinguishing variable (*e.g.* living with air conditioning versus not) and measuring OR. More targeted deployments may require finding such a factor that predicts similar disparity as the serosurvey approach.

For our practical examples, the model parameters are:

| Country | $\mathbf{S}^{+}$ | $\rho_H$ | $\log_{10}(\mathrm{OR})$ |
|---|---|---|---|
| Indonesia | 0.8806040 | 0.4668654 | 1.777455 |
| Malaysia | 0.5072371 | 0.1401441 | 31.251271 |
| Philippines | 0.8419634 | 0.3162632 | 26.265428 |
| Thailand | 0.7342112 | 0.2579016 | 28.575004 |
| Vietnam | 0.5870976 | 0.3116296 | 32.564970 |
| Peru | 0.7684385 | 0.2940011 | 1.566647 |

TABLE III. Fits to seroprevalence data.

For our sensitivity study, we used the following three by three grid for force of exposure and combined high risk fraction and disparity:

| Force of Exposure | Risk Fraction & Disparity: Low | Medium | High |
|---|---|---|---|
| Low | $\{0.5, 0.5, 2\}$ | $\{0.5, 0.3, 17.5\}$ | $\{0.5, 0.1, 33\}$ |
| Medium | $\{0.7, 0.5, 2\}$ | $\{0.7, 0.3, 17.5\}$ | $\{0.7, 0.1, 33\}$ |
| High | $\{0.9, 0.5, 2\}$ | $\{0.9, 0.3, 17.5\}$ | $\{0.9, 0.1, 33\}$ |

TABLE IV. Sensitivity categories, $\mathbf{S}^{+}, \rho_H, \log_{10}\mathrm{OR}$.

### IV. CONTEXT ECONOMICS

For the two practical examples in the main text, Peru and Malaysia, we use previously assumed cost data for $S$, $V$, and $T$ to set $\nu$ and $\tau$ [4].

The generally assumed cost of a vaccine regimen is 78 USD (21 USD per dose, for a three dose course, plus 15 USD for additional logistical costs), and test costs have been suggested at approximately 5 USD.

For Malaysia, if we assume the public payer perspective, the benefit of preventing a second-like infection is roughly $S = 86$ USD, therefore $\nu \approx .9$ and $\tau \approx .2$. For Peru, if we assume the public payer perspective, $S = 223$, $\nu \approx .3$ and $\tau \approx .1$.

In these two settings, we also estimated $\frac{F}{S}$; in Malaysia, it is $\frac{F}{S} \approx 0.2$ while in Peru $\frac{F}{S} \approx 0.4$. For our comparison to vaccination without testing, we assumed $\frac{F}{S} = 0.5$ as an upper limit.

[1] Maïna L'Azou, Annick Moureau, Elsa Sarti, Joshua Nealon, Betzana Zambrano, T Anh Wartel, Luis Villar, Maria RZ Capeding, and R Leon Ochiai. Symptomatic dengue in children in 10 Asian and Latin American countries. *N Engl J Med*, 374(12):1155–1166, 2016. doi:10.1056/NEJMoa1503877.

[2] Amy C. Morrison, Sharon L. Minnick, Claudio Rocha, Brett M. Forshey, Steven T. Stoddard, Arthur Getis, Dana A. Focks, Kevin L. Russell, James G. Olson, Patrick J. Blair, Douglas M. Watts, Moises Sihuincha, Thomas W. Scott, and Tadeusz J. Kochel. Epidemiology of dengue virus in Iquitos, Peru 1999 to 2005: Interepidemic and epidemic patterns of transmission. *PLoS Negl Trop Dis*, 4(5):1–17, May 2010. doi:10.1371/journal.pntd.0000670.

[3] Ankit Rohatgi. Webplotdigitizer, 2011-2019. URL `http://arohatgi.info/WebPlotDigitizer/`.

[4] Stefan Flasche, Mark Jit, Isabel Rodríguez-Barraquer, Laurent Coudeville, Mario Recker, Katia Koelle, George Milne, Thomas J Hladish, T Alex Perkins, Derek AT Cummings, et al. The long-term safety, public health impact, and cost-effectiveness of routine vaccination with a recombinant, live-attenuated dengue vaccine (Dengvaxia): a model comparison study. *PLoS Medicine*, 13(11):e1002181, 2016. doi:10.1371/journal.pmed.1002181.
